## Supplementary information for "Two long-axis dimensions of hippocampal-cortical integration support memory function across the adult lifespan"

### Contents

S. Figure 1. Explained variance of gradients ... 1

S. Figure 2. G3 in relation to hippocampal subfields ... 2

S. Figure 3. Connectopic mapping in an independent dataset ... 3

Stability of gradients across levels of spatial smoothing ... 4

S. Figure 4. Connectopic mapping on true and random data at varying levels of spatial smoothing ... 5

S. Figure 5. Selection of trend surface model (TSM) order ... 6

S. Figure 6. G2 and dopamine D1 receptor distribution ... 7

S. Table 1. Gradient topography as a predictor of episodic memory performance ... 8

Classification of older adults based on right-hemisphere G1 parameters ... 9

Replication of older classes in an independent sample ... 9

S. Figure 7. Classification of older adults based on gradient topography ... 10

References ... 11

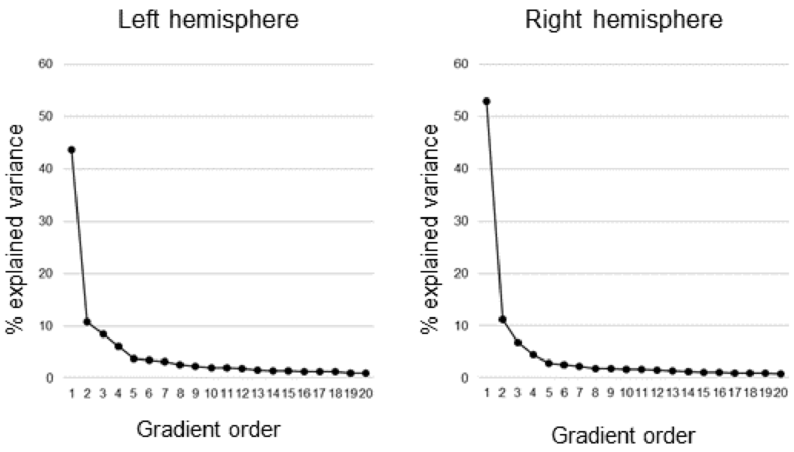


S. Figure 1. Explained variance of gradients. Connectopic mapping was used to compute 20 hippocampal gradients in each hemisphere on group level across the sample. The first-order gradient explained 44% and 53% of the variance in left and right hemispheres, respectively. The second-order gradient explained 11% in both hemispheres, and the third-order gradient 8% and 7% in left and right hemispheres.

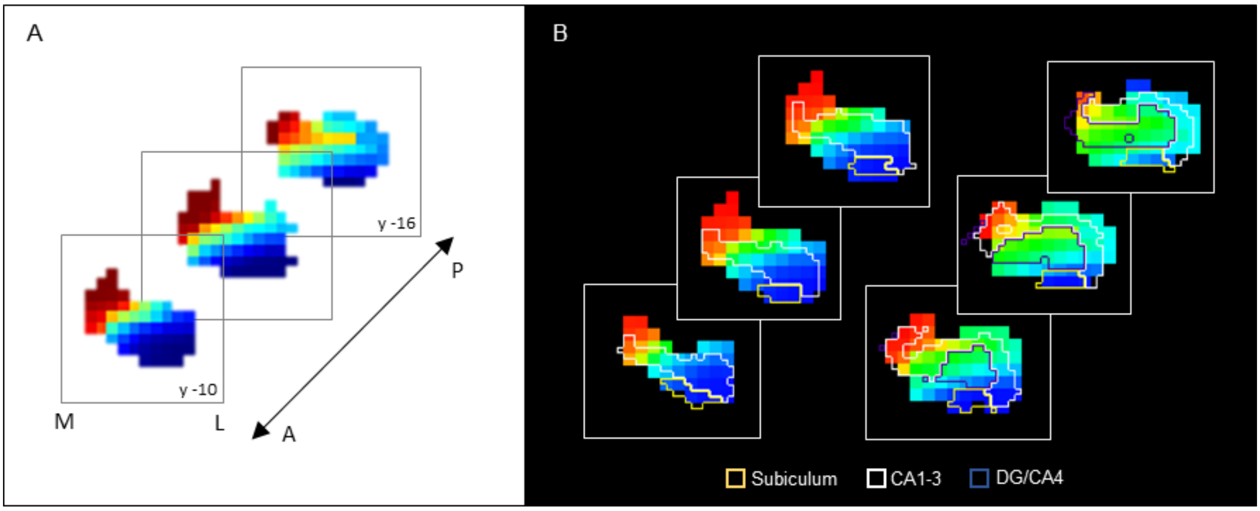


S. Figure 2. G3 in relation to hippocampal subfields. Coronal view of the third-order connectopic map, G3, within the anterior hippocampus, where medial-lateral variation was most evident. Hippocampal subfields are displayed as contours: subiculum (yellow), CA1-3 (white), DG/CA4 (blue).

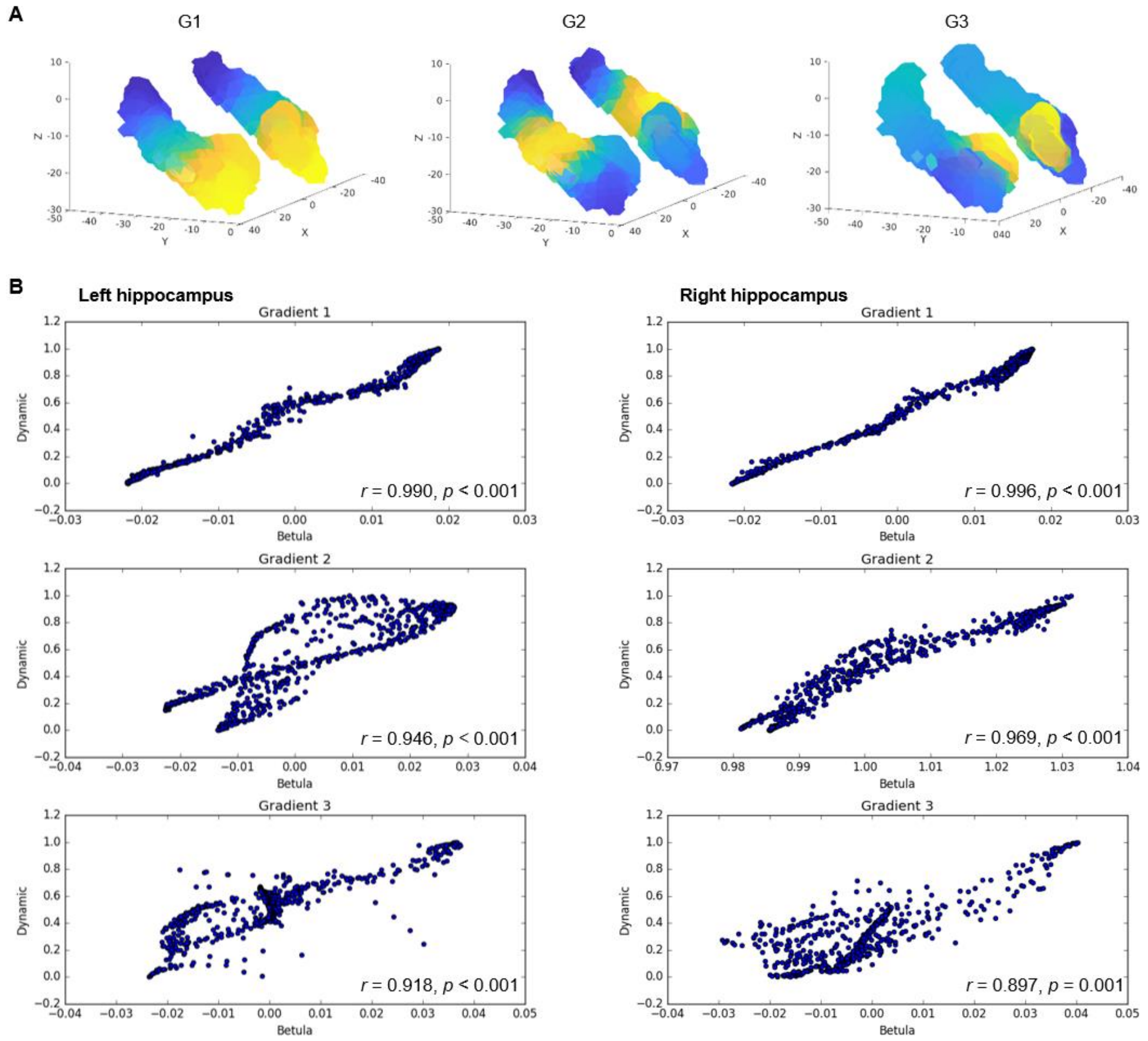


S. Figure 3. Connectopic mapping in an independent dataset. A) Gradients of hippocampal cortical connectivity were replicated in an independent sample of 224 adults (122 men/102 women; 29-85 years, mean age = 65.0±13.0) from the Betula project (Nilsson et al., 2004; Nyberg et al., 2020), such that a principal anterior-posterior gradient (G1) was followed by a second-order middle-to-anterior/posterior gradient (G2) and a third-order inferior-lateral to medial gradient (G3). B) Voxel-wise correlations between gradients in the Dynamic and Betula datasets.

### Stability of gradients across levels of spatial smoothing

Spatial smoothing is commonly implemented during preprocessing of functional MRI data. Benefits of smoothing include enhancing the signal-to-noise ratio and reducing effects of anatomical variability between subjects (Molloy et al., 2014; Wu et al., 2011). On the other hand, smoothing is associated with greater spatial autocorrelation. In a recent study, Watson and Andrews (2023) suggested that spatial autocorrelation induced by smoothing during preprocessing is sufficient for detecting functional gradients through connectopic mapping, even in artificially generated random fMRI time series (Watson & Andrews, 2023). To determine the stability of our hippocampal gradients, in light of smoothing as a potential confound, we ran connectopic mapping on both true and random data at varying levels of spatial smoothing (i.e., no smoothing, 0.5mm, and 6.0mm).

Random fMRI data were generated according to the methods specified by Watson and Andrews (2023). Briefly, we synthesized Gaussian white noise matched in mean and variance to the real fMRI data in native space. The random time series therefore approximate the signal amplitude and variation of the true data, without a coherent spatial or temporal correlation structure. The random fMRI data were then normalized to MNI space with varying levels of smoothing. Normalization for both true and random data was performed using DARTEL (Ashburner, 2007) implemented in SPM12. This method projects the location of voxels to a template, effectively preserving the tissue count, but is known to introduce aliasing artifacts if the original data is at a similar or lower resolution than the deformation fields. The effects of aliasing artifacts are typically minimized by smoothing the data during the normalization procedure. For non-smoothed data, missing voxels along the X and Y dimensions were replaced by interpolating neighboring voxels along the Z-dimension. This effectively removes aliasing artifacts by resampling a single voxel along the projected dimension, avoiding spurious autocorrelations otherwise produced by smoothing.

Our results support the detection of expected hippocampal gradients in true data independent of smoothing degree (S. Figure 4A-B). Age-related variance was captured in both left and right G1 at all levels of smoothing, whereas additionally visible across G2 and G3 from a minimum level of 0.5mm smoothing (S. Figure 4C). In contrast, cmaps derived from random data showed poor consistency with expected gradients (S. Figure 4B) and could not capture individual differences (S. Figure 4C). In sum, the hippocampal gradients in our work are highly stable in their spatial layout, and in their ability to capture inter-individual differences, across levels of spatial smoothing. Observations in random data indicate that spatial smoothing is not sufficient to produce meaningful gradients of functional variance in the hippocampus.

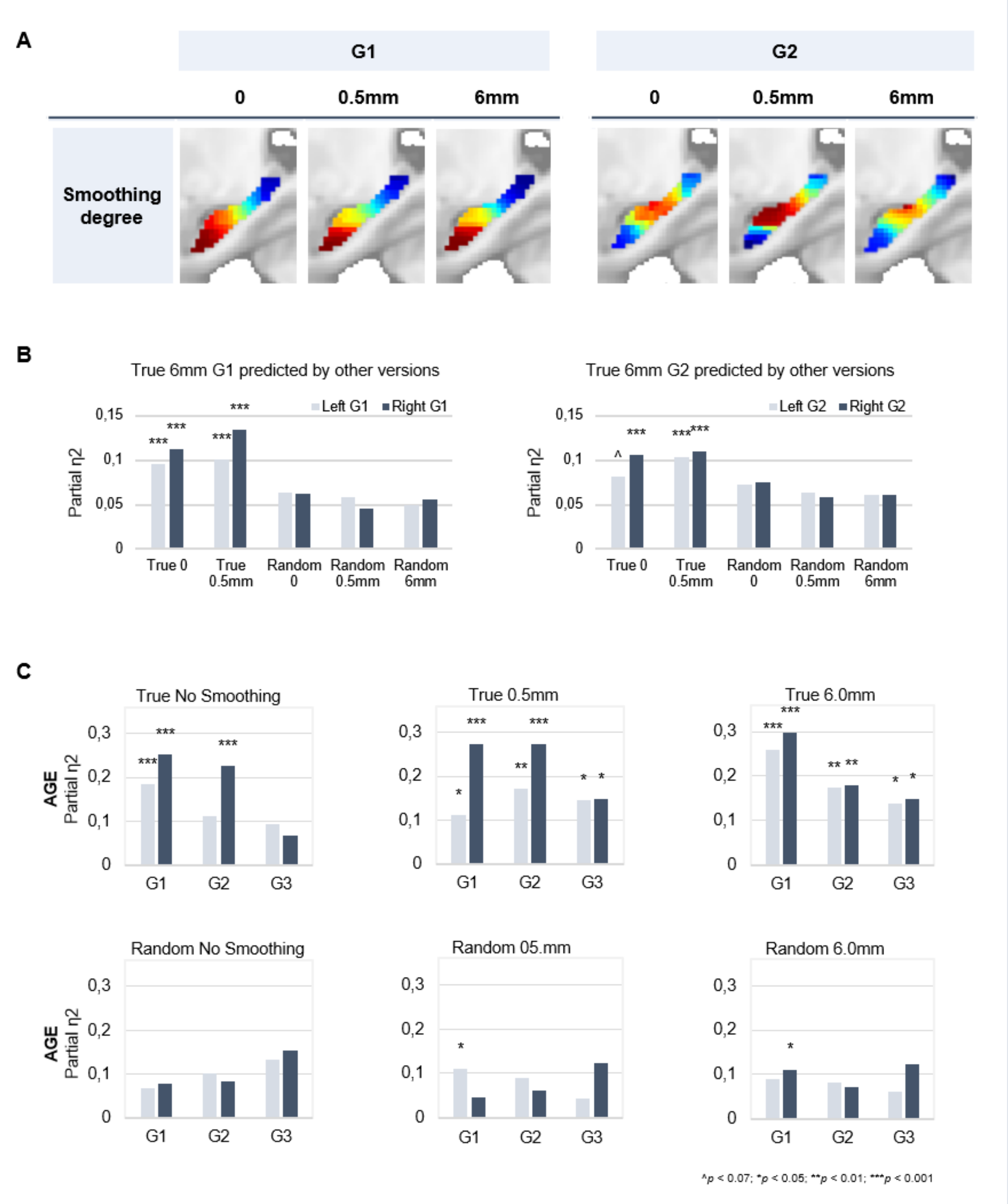


S. Figure 4. Connectopic mapping on true and random data at varying levels of spatial smoothing. A) G1 (left panel) and G2 (right panel) connectopic maps derived from resting-state fMRI data after different degrees of spatial smoothing. B) Bars represent effect sizes from multivariate GLMs with TSM parameters of the G1 (left) and G2 (right) connectopic maps derived from 6.0mm smoothed data (presented in the main text) as dependent variables, predicted by TSM parameters of connectopic maps based on i) true data not smoothed; ii) true 0.5mm smoothed data; iii) random data not smoothed; iv) random 0.5mm smoothed data; v) random 6.0mm smoothed data. C) Effects of age on the topographic characteristics of gradients (i.e., TSM parameters) for true and random versions across levels of spatial smoothing.

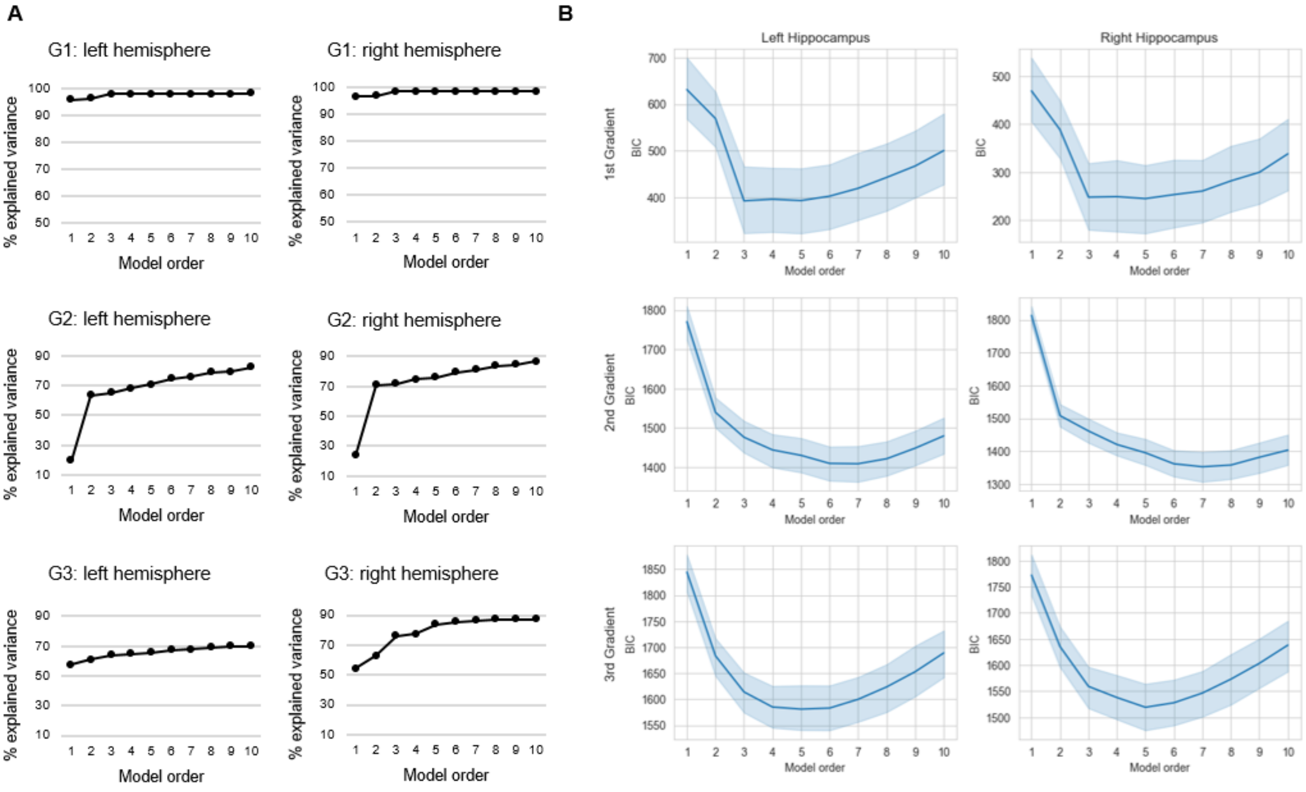


S. Figure 5. Selection of trend surface model (TSM) order. A) The % explained variance in group-level connectopic maps by TSM models. B) Average values of the Bayesian Information Criterion (BIC) across participants are plotted against trend surface model orders. Shaded areas represent the 95% confidence interval.

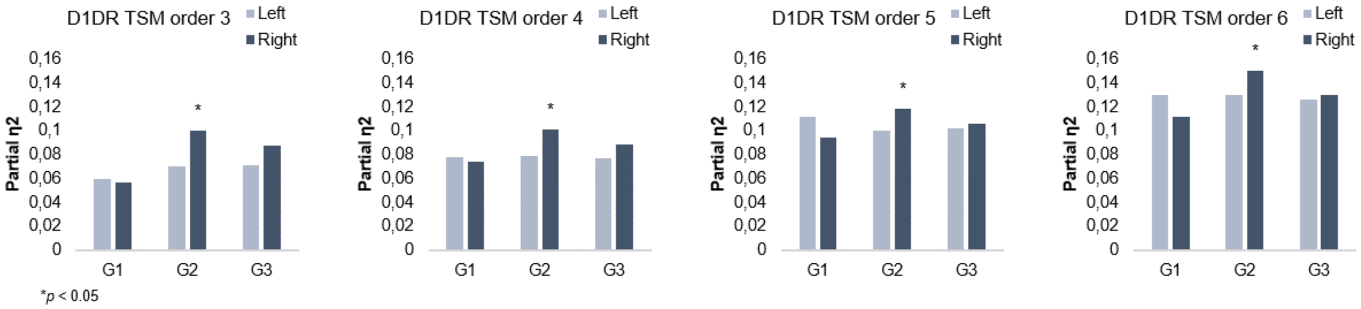


S. Figure 6. G2 and dopamine D1 receptor distribution. Individual differences in D1DR topography predict G2 topography in the right hemisphere across TSM model orders.

S. Table 1. Gradient topography as a predictor of episodic memory performance

| | Full sample | | Young (20-40y) | | Middle-aged (40-60y) | | Older (60-79y) | |
|---|---|---|---|---|---|---|---|---|
| | $\Delta R^2$ | F | $\Delta R^2$ | F | $\Delta R^2$ | F | $\Delta R^2$ | F |
| Model 1[1] | 0.287 | 20.92*** | 0.003 | 0.058 | 0.131 | 2.567^ | 0.163 | 2.915* |
| *Left HC* | | | | | | | | |
| G1 | 0.029 | 0.695 | 0.357 | 2.672* | 0.076 | 0.450 | 0.156 | 0.916 |
| G2 | 0.096 | 1.842* | 0.191 | 1.098 | 0.264 | 1.246 | 0.215 | 0.921 |
| G3 | 0.027 | 0.502 | 0.236 | 1.755 | 0.261 | 1.462 | 0.256 | 1.219 |
| *Right HC* | | | | | | | | |
| G1 | 0.026 | 0.621 | 0.150 | 0.810 | 0.128 | 0.804 | 0.169 | 0.987 |
| G2 | 0.056 | 1.013 | 0.328 | 1.542 | 0.221 | 1.066 | 0.151 | 0.566 |
| G3 | 0.036 | 0.633 | 0.120 | 0.432 | 0.288 | 1.865 | 0.276 | 1.070 |

[1]M1 = age, sex, mean frame-wise displacement; ^$p < 0.07$; *$p < 0.05$; **$p < 0.01$; ***$p < 0.001$

### Classification of older adults based on right-hemisphere G1 parameters

Classification of older adults based on right-hemisphere G1 TSM parameters yielded a two-class solution, by definition, these two groups differed from each other in terms of right-hemisphere G1 characteristics ($F_{(9,37)} = 10.886$, $p < 0.001$, partial $\eta^2 = 0.726$), with a difference between groups also evident in the left hemisphere ($F_{(9,37)} = 2.248$, $p = 0.040$, partial $\eta^2 = 0.353$). The two sub groups primarily showed a youth-like (vs. young: $F_{(9,73)} = 1.428$, $p = 0.192$, partial $\eta^2 = 0.150$) vs. aged (vs. young: $F_{(9,70)} = 8.574$, $p < 0.001$, partial $\eta^2 = 0.524$) gradient profile in terms of right-hemisphere G1 parameters (S. Figure 6A), as such, the classification did not extend across the other gradients like the one based on left-hemisphere G1 parameters. The two sub groups did not significantly differ in terms of age (aged: 70.0±5.8; youth-like: 68.5±5.0; $t = 1.030$, $p = 0.308$), sex (aged: 13 men/10 women; youth-like: 13 men/13 women; $X^2 = 0.208$, $p = 0.648$), nor hippocampal gray matter volume (left hemisphere: aged: 4345.4±448.4; youth-like: 4173.1±269.0; $t = 1.614$, $p = 0.114$; right hemisphere: aged: 3981.2±428.2; youth-like: 3874.4±426.9; $t = 0.872$, $p = 0.388$). Sub groups identified based on right-hemisphere G1 topography did not significantly differ in episodic memory performance (the composite episodic measure used across the sample: aged: 43.9±5.01; youth-like: 45.6±6.0; $t = 1.093$, $p = 0.280$; word recall sub test: aged: 42.0±5.2; youth-like: 44.9±7.8; $t = 1.545$, $p = 0.129$).

### Replication of older classes in an independent sample

The ability of left G1 topography to inform classification of older adults into mnemonically distinct subgroups was replicated in the Betula sample (class 1: n = 60; class 2: n = 99; $F(9,149) = 35.993$, $p < 0.001$, partial $\eta 2 = 0.685$), S. Figure 7C. A difference in gradient topography between classes was also evident for G1 in the right hemisphere ($F(9,149) = 2.134$, $p = 0.030$, partial $\eta 2 = 0.114$), but was more limited across subsequent gradients in both hemispheres. The smaller class was, in relation to young adults, exhibiting an aged G1 ($F(9,63) = 4.166$, $p < 0.001$, partial $\eta 2 = 0.373$) in contrast to the bigger class displaying a more youth-like G1 ($F(9,102) = 1.345$, $p = 0.223$, partial $\eta 2 = 0.106$). Importantly, we observed a trend-level difference in word recall performance between subgroups ($t(157) = 1.585$, $p = 0.058$, 1-tailed), such that youth-like older adults displayed superior memory performance.

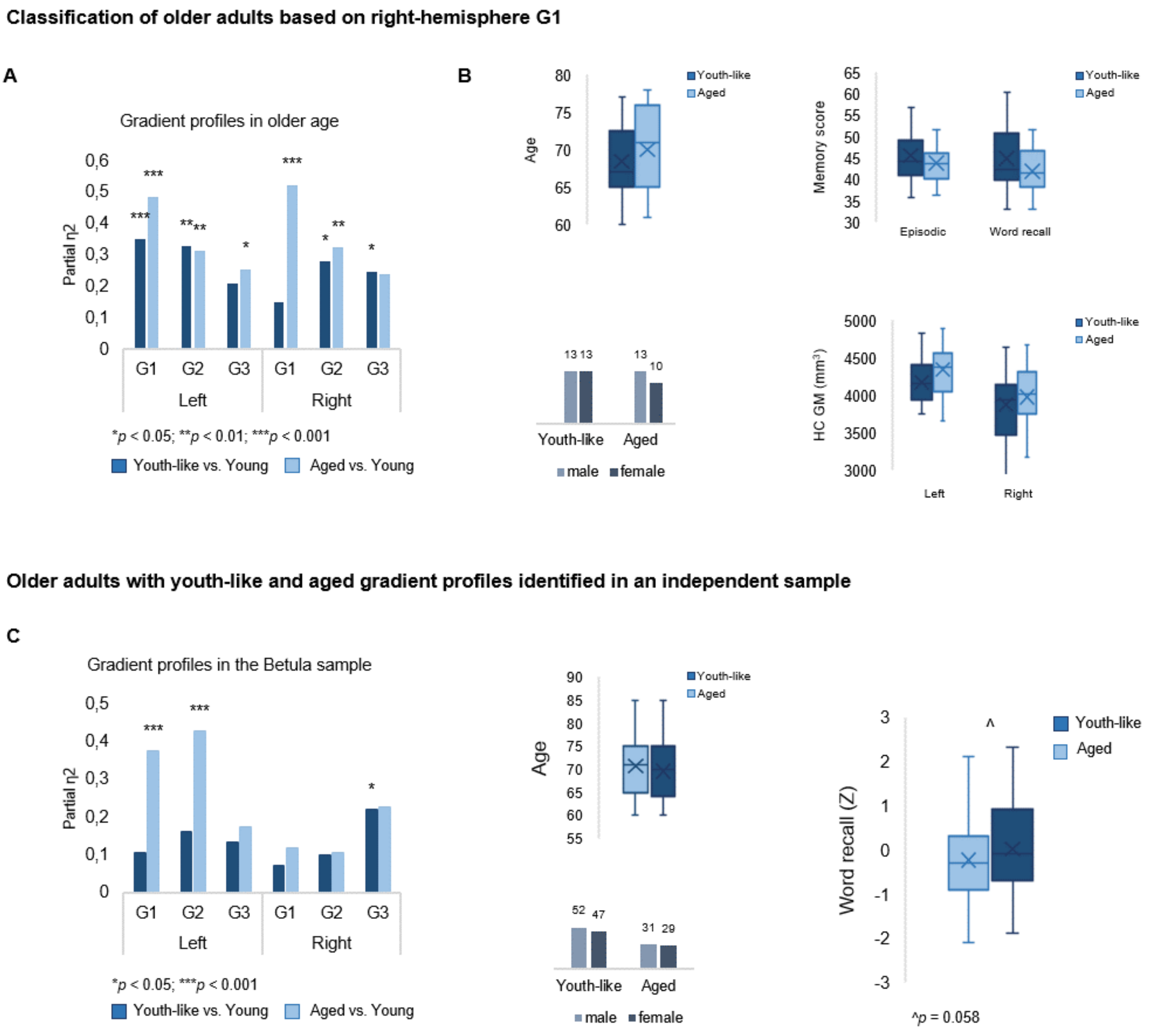


S. Figure 7. Classification of older adults based on gradient topography. A) Classification of older adults in DyNAMiC based on right-hemisphere G1 parameters. The first group (n=23) displayed right G1 characteristics significantly different from those in young adults, whereas the second group (n=26) displayed right G1 characteristics more similar to those in young adults. B) Older sub groups were comparable in terms of age, sex, hippocampal gray matter (GM) volume, and memory performance. C) Classification of older adults in the independent sample, Betula, based on left-hemisphere G1. A subgroup with a youth-like G1 displayed higher episodic memory performance.

### References

Ashburner, J. (2007). A fast diffeomorphic image registration algorithm. *NeuroImage*, *38*(1), 95–113. https://doi.org/10.1016/j.neuroimage.2007.07.007

Molloy, E. K., Meyerand, M. E., & Birn, R. M. (2014). The influence of spatial resolution and smoothing on the detectability of resting-state and task fMRI. *NeuroImage*, *86*, 221–230. https://doi.org/10.1016/j.neuroimage.2013.09.001

Nilsson, L.-G., Adolfsson, R., Bäckman, L., Frias, C. M. de, Molander, B., & Nyberg, L. (2004). Betula: A Prospective Cohort Study on Memory, Health and Aging. *Aging, Neuropsychology, and Cognition*, *11*(2–3), 134–148. https://doi.org/10.1080/13825580490511026

Nyberg, L., Boraxbekk, C.-J., Sörman, D. E., Hansson, P., Herlitz, A., Kauppi, K., Ljungberg, J. K., Lövheim, H., Lundquist, A., Adolfsson, A. N., Oudin, A., Pudas, S., Rönnlund, M., Stiernstedt, M., Sundström, A., & Adolfsson, R. (2020). Biological and environmental predictors of heterogeneity in neurocognitive ageing: Evidence from Betula and other longitudinal studies. *Ageing Research Reviews*, *64*, 101184. https://doi.org/10.1016/j.arr.2020.101184

Watson, D. M., & Andrews, T. J. (2023). Connectopic mapping techniques do not reflect functional gradients in the brain. *NeuroImage*, 120228. https://doi.org/10.1016/j.neuroimage.2023.120228

Wu, C. W., Chen, C.-L., Liu, P.-Y., Chao, Y.-P., Biswal, B. B., & Lin, C.-P. (2011). Empirical Evaluations of Slice-Timing, Smoothing, and Normalization Effects in Seed-Based, Resting-State Functional Magnetic Resonance Imaging Analyses. *Brain Connectivity*, *1*(5), 401–410. https://doi.org/10.1089/brain.2011.0018
